# Membrane voltage and connexin expression work together to enhance tumor growth and metastasis in cancer

**DOI:** 10.64898/2026.09.02.748840

**Authors:** Joel Grodstein, Juanita Mathews, Michael Levin

## Abstract

There is strong evidence of tumors manipulating their resting membrane potential (*V*_mem)_. While most fully-differentiated cells have a *V*_mem_ of roughly -70mV, tumor cells are generally depolarized, with *V*_mem_ ≈-30mV, which more closely resembles the *V*_mem_ of stem cells. This is often believed to serve the purpose of accelerating the cell cycle and hence advantaging tumor proliferation. But when the tumor becomes invasive, its cells sometimes revert to a hyperpolarized *V*_mem_ with no obvious reason why. Separately, it is well accepted that solid tumors that are not yet invasive greatly underexpress connexins relative to healthy tissue; connexins, for our purpose, form gap junctions (GJs), small connecting tubes between nearby cells. Tumors that are invasive, by contrast, overexpress connexins. There is very little explanation for the paradox that connexins are first underexpressed and then overexpressed. However, it has long been known that *V*_mem_ electrically gates GJs; specifically, that homotypic GJs conduct best when the two cells they connect have a similar *V*_mem_. Our in-silico model results explain this phenomenon, showing that when considered together, tumors’ electrical and connexin-expression behaviors form a unified and effective strategy to control communication between the tumor and its healthy neighbor cells. This has implications for the emerging field of cancer bioelectrics, potentially leading to more precisely-targeted therapies.

## Introduction

The bioelectric state of cells, defined as the resting potential across their plasma membrane, *V*_mem_, is increasingly recognized as an important driver of cell behavior, morphogenesis, and health/disease outcomes [1–4]. This is especially relevant to cancer, because bioelectric networks are a powerful endogenous medium for binding individual cells toward organ-level set points and away from unicellular defections such as aging and neoplasia [5–7]. Key components of these circuits include ion channels [8, 9] and the electrical synapses known as gap junctions, made of connexin proteins [10].

There is strong evidence (reviewed in [8, 11–13]) of tumors changing their resting membrane potentials:

- Tumor cells are generally depolarized, with a *V*_mem_ of roughly -30mV, more closely resembling stem cells than the *V*_mem_ of fully differentiated cells.
- When the tumor becomes invasive, most experiments show it remaining depolarized (reviewed in [14]), but others show it reverting to a hyperpolarized *V*_mem_ [15].

Separately, there is strong evidence (reviewed in [16–18]) of altered connexin expression in tumors. Connexins are part of several structures relevant to tumors [17], notably gap junctions (GJs). A GJ is a small conduit between neighboring cells that selectively conducts molecules between the two cells. GJ density is decreased (relative to healthy tissue) in proliferating solid tumors, and restoration of GJ communication can slow tumor proliferation [18]. Some chemotherapy agents act by encouraging GJ formation in tumors [16], and transfecting GJ DNA into tumor cells can rescue the tumor [16]. Tumors that are invasive, by contrast, overexpress GJs [16–18].

While the data above are well documented, hypotheses as to the *why* are less clear. The initial depolarization has been seen as a driver of dedifferentiation and of accelerated replication [8]; indeed, forcibly preventing depolarization has prevented dedifferentiation and replication in some glioblastoma lines [19]. But there are few hypotheses for the connection between *V*_mem_ and metastasis. Ion-channel-expression changes (which typically change *V*_mem_) have been suggested to be related to motility in metastasis [13, 20], and rhythmic ion-channel changes can cause rhythmic cell-volume changes that aid motility[20]. But this does not explain why different tumors have different *V*_mem_ behavior during metastasis, or even attempt to explain why connexins are first underexpressed and then overexpressed.

The molecular pathways for connexin dysregulation in cancer are not yet fully understood [18]. It is not even clear whether the decreased GJ density in proliferating tumors is always a direct consequence of decreased connexin expression; it may be because connexins that are expressed do not successfully travel to the cell membrane to become part of GJs [18, 21, 22].

It is not obvious to see how both GJ underexpression and also overexpression could each be adaptive behavior for tumors, and how both depolarization and hyperpolarization could be adaptive. Presumably [16] the strategy that is best for proliferation may not be best for metastasis – but how exactly?

We propose a single hypothesis that explains both GJ initial underexpression and subsequent overexpression, ties in the existing data on *V*_mem_, and explains why one behavior is adaptive for proliferation and a nearly-opposite behavior is appropriate for invasion. This is important because it may help break the logjam preventing cancer therapies that target connexin misexpression. Since connexins are initially underexpressed and later overexpressed, we can only rarely identify them as oncogenes. Furthermore, while there is currently no obvious way to use them as chemotherapy targets, being able to target connexins only in some voltage states may help to more precisely target them. Finally, understanding both *V*_mem_ and connexin changes as a means of controlling communication may help to focus research on exactly what messages are being sent, eventually providing still more targets.

Our hypothesis is based on the observation, known since the 1980s ([23], and reviewed in [24]) that GJs are voltage sensitive. Thus, the tumor *V*_mem_ changes *must* affect GJ conductivity. One of the simplest types of GJs is a *homotypic* GJ, which is made up of a connection using the same type of connexin on both joined cells and which conducts best when the two cells that it connects have roughly the same *V*_mem_ [24, 25] (Figure 1). It doesn’t matter what that *V*_mem_ is, as long as it is the same in both cells; i.e., homotypic-GJ conductance depends primarily on |Δ*V*_mem_| between the two cells that it connects, rather than *V*_mem_ of either individual cell.

**Figure 1:**
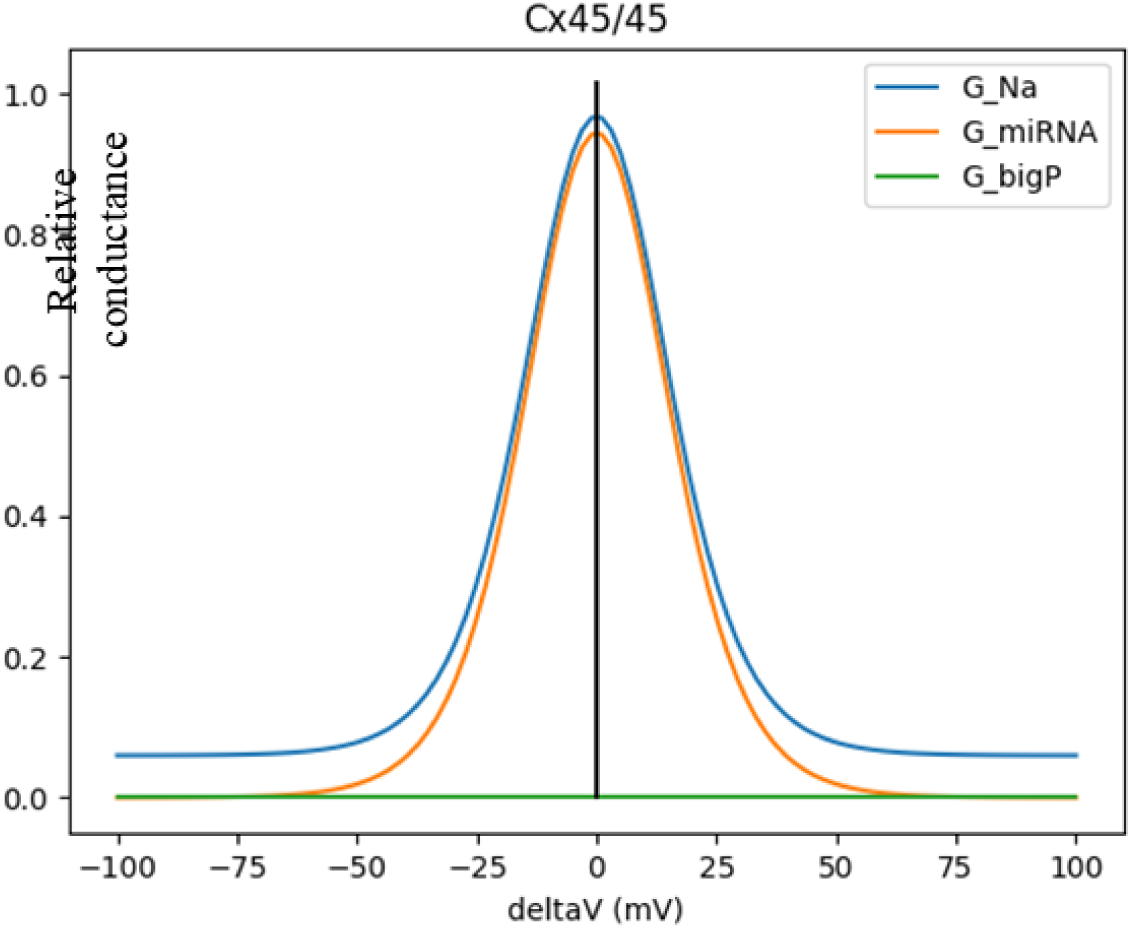
*GJ conductance varies with ΔVmem.* Graph of conductance vs. ΔV_mem f_or a homotypic GJ. Homotypic means that both hemichannels are built from the same connexin (in this case, Cx45). Any individual GJ is either on or off; the overall conductance is determined by the fraction of GJs that are on.

When the cells of a tumor all depolarize, it does not change the Δ*V*_mem_ across GJs within the tumor. However, it does change the Δ*V*_mem_ of the border GJs – i.e., those GJs that connect a depolarized (tumor) cell to a hyperpolarized (healthy) cell. Thus, one effect of *V*_mem_ depolarization is to reduce communication between the tumor and the surrounding healthy tissue.

Why would communication be so important? The ability of healthy surrounding tissue to normalize some malignancies has been suspected since [26–28], noted in [17, 18], and reviewed in context of the somatic mutation theory in [7, 29]. Cancer biology encompasses a wide array of tumor strategies and it is difficult to generalize the types of messages that healthy tissue sends, but we only care that they exist and can be transmitted through GJs.

### Summary of our hypothesis

When a small tumor starts to proliferate, the surrounding healthy tissue will try to normalize the tumor by sending messages. Since the tumor is still small, it may have a hard time resisting these normalizing messages – and so its best strategy may be to cut communication with the surrounding healthy tissue (likely an ancient response of unicellular organisms to prevent bystander-toxicity effects and exploitation). First, it underexpresses connexins, which reduces gap-junctional communication with the surrounding tissue. Second, it depolarizes, which creates a large Δ*V*_mem_ across the border GJs that connect the tumor to nearby healthy tissue. This lowers the conductance of these border GJs, further disconnecting the tumor from its surrounding healthy tissue. However, since all of the tumor cells have the same depolarized *V*_mem_, the intra-tumor GJs still have Δ*V*_mem_=0 and are thus still fully conducting, leaving intra-tumor communication relatively intact (other than the decreased GJ density). The effect of this is that the tumor cells thus communicate well with each other and very little with the healthy surrounding tissue. We thus hypothesize that, in addition to any function controlling differentiation and cell division, the depolarization is also about controlling communication.

If communication through GJs is an important part of the healthy body’s strategy for stopping tumor proliferation, then minimizing that communication becomes essential for a tumor’s survival, explaining why depolarizing *V*_mem_ would be part of a tumor’s growth strategy. This also explains why the tumor underexpresses connexins; doing so helps it to ignore signals sent to it that might otherwise restore homeostasis.

But why, then, would a tumor reverse course and overexpress connexins during invasion and metastasis? We hypothesize that it is still about communication. Several authors ([30–32], and summarized in [18]) have noted, both in vitro and in vivo, that the GJ overexpression in invasive tumors is especially marked near the tumor’s border with healthy tissue, and have thus hypothesized that the tumor uses GJs to send metabolites to alter the neighboring healthy tissue and thus ease invasion.

We agree; and will show, both through simulation and analytically, that GJ overexpression essentially turns the tumor into a syncytium. A small tumor had to isolate itself to prevent being normalized; but a larger tumor, by overexpressing GJs and thus acting as a single large, dominant cell, can send signals strong enough to override the body’s signals to nearby healthy cells, subverting those nearby cells to its own purpose (similar effects at another scale have been observed in different sized sets of embryos resisting teratogenic influence [33]). The change in GJ expression thus marks the tumor’s switch from isolation to proselytization. When the tumor’s goal was isolation, GJ underexpression was adaptive; but when its goal is invasion, overexpression becomes a far better strategy.

But GJ overexpression is only half of a proselytization strategy. As noted above, most GJs are voltage gated. If the tumor remained depolarized, then the border GJs would still remain closed and thus isolate the tumor from healthy tissue, preventing the tumor from effectively transferring signaling molecules. The literature is mixed on whether invasive tumors remain depolarized or re-hyperpolarize [12, 13, 15, 20]. We will propose mechanisms by which either can aid its proselytization.

## Methods

We now discuss our detailed simulation model. Each part of our hypothesis will be tested by numerical simulations. Our simulator uses numerical integration to simulate multiple cells interconnected by GJs, with each cell containing individual ion channels, Na-K-ATPase pumps and arbitrary GRNs. The ion channels use the Goldman-Hodgkins-Katz model [34–36]; the pumps use the thermodynamically-driven model from [35, 36].

Our goal is to simulate genetic circuits holding tumor-related state. But cancer is wide and varied, and different tumors may transfer different metabolites through GJs to affect different pathways. We decided to focus on miRNA passing through GJs. While GJs are not known to pass proteins between cells [24], miRNA passing through GJs has been demonstrated in both the GJ [25, 37] and cancer [17] literature.

Over half of all human genes are targeted by some form of miRNA [38], and miRNA can specifically target connexin coding [17]. MiRNA has been shown to regulate cell growth and proliferation (reviewed in [39, 40]) and apoptosis (reviewed in [41]), again making it a logical choice to affect cancer.

MiRNA is underexpressed in tumors [42], which has been interpreted [43] as the tumor wanting to avoid the regulation of its own behavior by miRNA. It has been shown to be highly differentially expressed in gliomas and to inhibit glioma motility and invasiveness [44]. Transfer of miR-5096 derived from glioma cells to astrocytes through GJs promotes pro-invasive behavior [45]; similar results apply for miR-19b [46]. Establishment of GJs between normal neural cells and tumor cells can attenuate cell proliferation due to miR124-3p transfer [47]. In melanoma, hypoxic tumor cells transfer miR-192-5p to dendritic cells and tumor-associated T-lymphocytes via Cx40, suppressing the cytotoxic activity of T-lymphocytes [48].

We have thus chosen to use a miRNA circuit as our simulation model; specifically, the ubiquitous EMT switch [49, 50]. All carcinomas must switch from an E (epithelial) to M (mesenchymal) phenotype in order to metastasize, and EMT pathways are well studied. We use the pathways and model from [50], shown in Figure 2. The circuit holds state due to a cross-coupled loop between ZEB1 and miR200.

**Figure 2:**
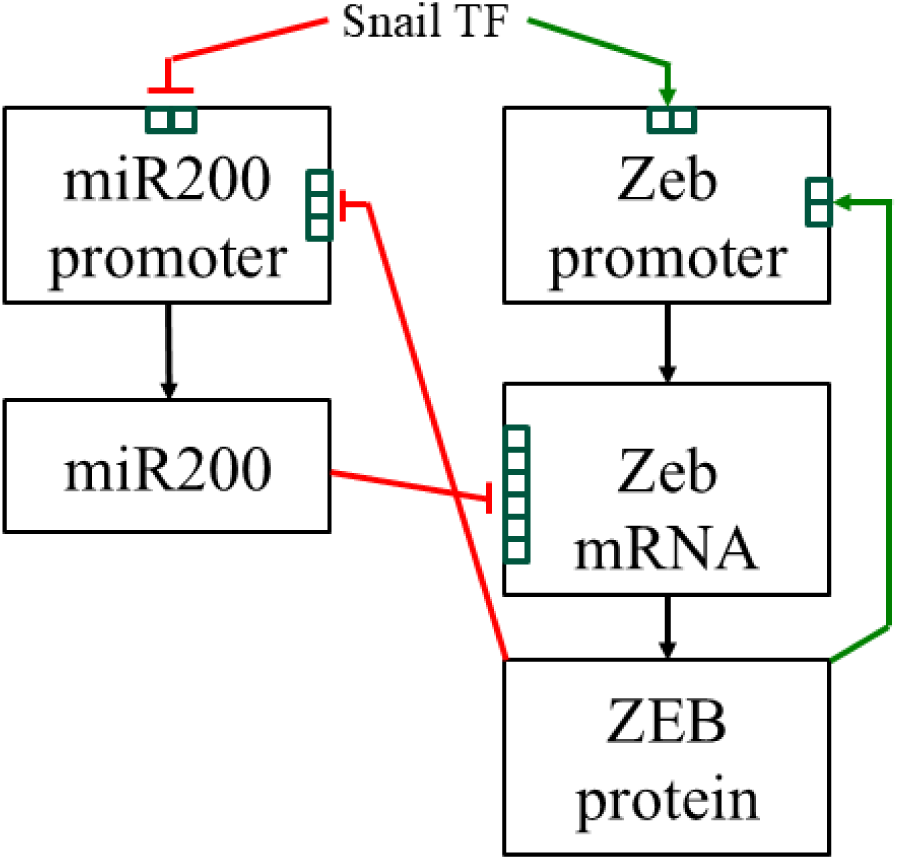
*GRN-level diagram of the EMT switch.* The two top boxes are the promoters for miR200 and Zeb; each of their small interior squares represents a promoter or repressor binding site. Green arrows represent activation and red bars represent repression. The two middle squares represent RNA. The six small squares in the Zeb mRNA box represent repression sites for cooperative binding of miR200 repressing the translation of Zeb mRNA to ZEB protein. Bistability is created by cross-repression: miR200 represses ZEB translation, and ZEB represses miR200 transcription. ZEB also has a self-loop, promoting its own transcription. The external transcription factor Snail can bias the circuit towards E, M or even a third hybrid state.

Each cell will contain an instance of the circuit shown in Figure 2, generating the miRNA miR200 (an E marker) and the protein ZEB (an M marker). Following the modeling in [49], we use modified Hill models for the activators and repressors, where 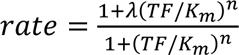 Values of λ>1 correspond to activation; 0< λ<1 implement repression. *K*_m_ is the Michaelis concentration, at which the output rate is half-maximum; *n* is the Hill-model cooperativity, and *TF* is the concentration of the input transcription factor (Snail for both promoters). We assume that the chemical bonding of miR200 to Zeb occurs substantially faster than transcription or translation, and can thus be treated as always at steady state.

The EMT switch is believed to support not only the E and M states, but also a hybrid H state that is highly invasive [49]. To simplify our simulations, we have chosen [Snail] to bias the circuit into a region where only E and M are stable. Our simulations show that the transfer of miR200 between cells through GJs can flip a cell from E to M or vice versa. This is what we simulate for the messages sent – the flipping of cells between E and M state. A tumor cell (in the M state) will be considered normalized if its state is flipped to E, and vice versa.

We next elaborate how we model miRNA passing through GJs via electrophoresis. Even though an isolated cell would then have only two stable states (E and M), when two cells of opposite state are interconnected by GJs the system becomes more complex. Figure 3a shows two cells connected by a GJ. We model the diffusion flux through GJs with Fick’s Law as 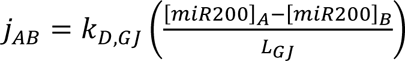, where *j*_AB_ is the flux from cell *A* to cell *B* in mol/m^2s;^ *k*_D,GJ_ is the diffusion constant for miR200 through the GJ, in m^2/s^; *L*_GJ_ is the length of the GJ in m; and the subscript “A” (“B”) denotes concentrations in cell A (cell B). If we define *G_GJ_* = 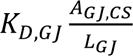 as a gap-junctional conductance in m^3/s^, where *A*_GJ,CS_ is the cross-sectional area of GJs between cells *A* and *B*, in m^2,^ then Fick’s Law becomes more simply *i_AB_* = *j_AB_A_GJ_*_,*CS*_ = *G_GJ_*([*miR*200]*_A_* − [*miR*200]*_B_*), where *i*_AB_ is the GJ current from *A* to *B* in mol/s.

**Figure 3.**
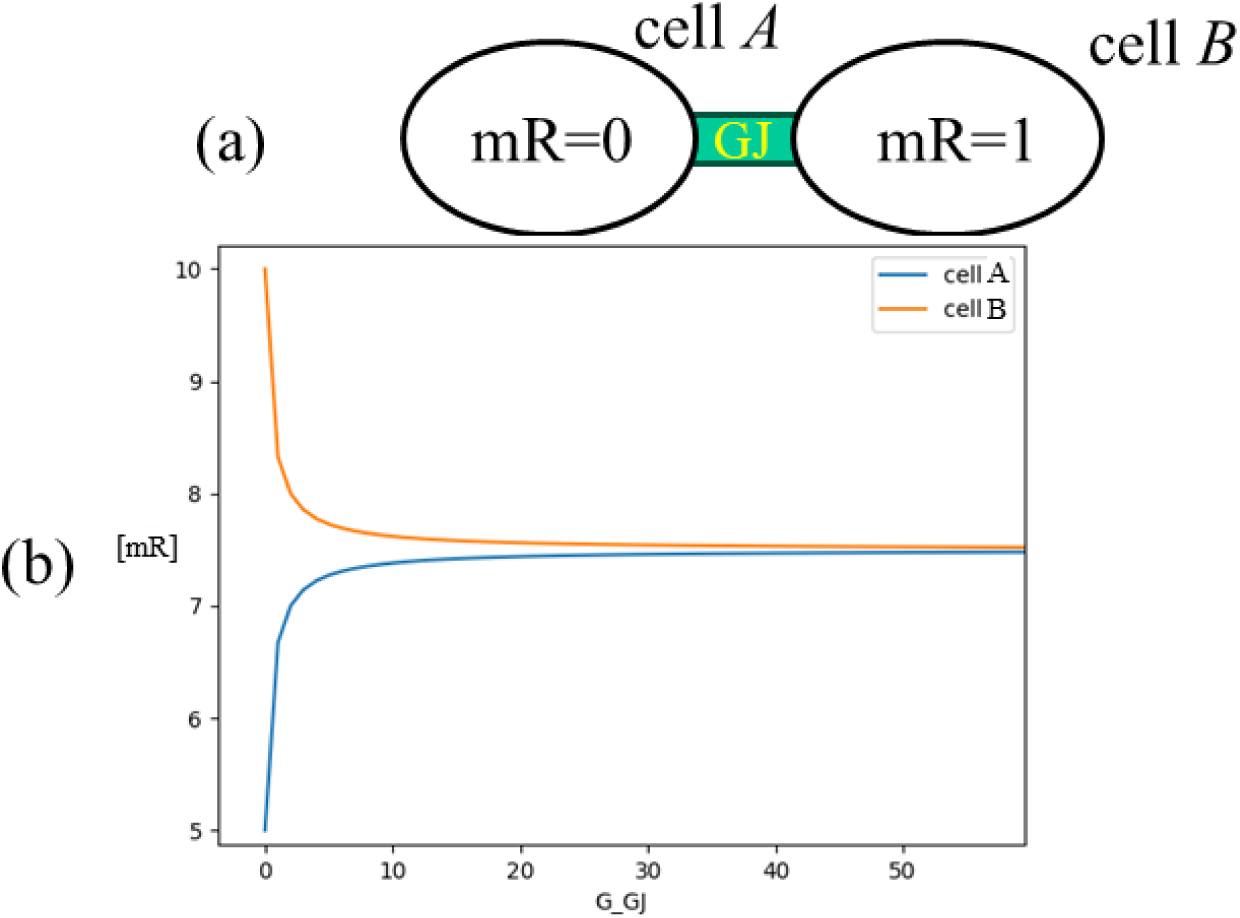
*two cells with opposite state connected by a GJ.* (a) Two cells connected by GJs. mR=0 and mR=1 denote the concentration of the relevant miRNA in each cell (with opposite state) before interconnection with the GJ. (b) Steady-state [mR] in each cell after the GJ is present. At t=0, cell A had [mR]=5 and cell B had [mR]=10. Then the GJ was added to the simulation. The x axis is not time, since these are steady-state results; it is *G*_GJ._

Any charged species is subject to drift currents. Our communicating miRNA has a large negative charge, and GJs also conduct small charged ions (e.g., Na, K and Cl). Fick’s Law then becomes the Nernst-Planck equation by adding drift, with mobility *μ* derived from *k*_D,GJ_ by the Einstein relation [51]. We also add voltage dependent GJ gating by the equation *G_GJ_* = 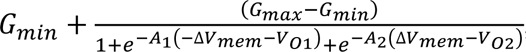, where *G*_min a_nd *G*_max_ are the absolute min/max conductances of the GJ, *V*_O1_ and *V*_O2_ control the width of the passband, and *A*_1_ and *A*_2_ control the steepness of the falloff [52].

However, modeling the flow of miRNA through GJs has two complexities. First, a GJ is more complex than a simple gated tubule. It is made of two hemichannels, each contributed by one of the two cells connected by the GJ. Each hemichannel then consists of two gates in series, typically denoted the “fast” and “slow” gate [24]. In principle, both gates can be controlled both by voltage and chemically [53]. The slow gate can close completely, blocking all ion passage – but is typically controlled only chemically, and is thus not part of our model. The fast gate typically closes electrically, but does not extend across the entire GJ pore. Thus, the fast gate, even when “fully” closed, does not fully block the flow of small ions. This is why the main blue curve in Figure 1 does not drop down to *G*_GJ_=0. However, closure of the fast gate does fully block larger ions [25, 37]. The experiments in the GJ literature use dye molecules to show fast-gate closure; we are not aware of any experiments using miRNA. However, since the miRNA diameter is nearly as wide as the entire GJ pore diameter, it seems very likely that they are fully blocked (Figure 1, orange curve).

There is a second, separate effect to consider. MiRNA’s effective charge is much smaller than its actual charge. Every nucleotide has a phosphate group, each of which carries a -1 charge, thus giving a 22-nucleotide miRNA a charge of -22. However, the cytoplasm is replete with mobile small ions; Manning theory [54, 55 ch 14] predicts that roughly 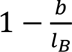 of the phosphate groups will bind so tightly to mobile cations as to essentially move as a unit (and are then called *counterions*), where *b* is the miRNA charge spacing (roughly .6nm) and *l*_B_ is the Bjerrum length (roughly .7nm) giving a remaining charge of -22*.6/.7≈-18.8.

Next, the miRNA molecule will electrostatically attract a cloud of mobile cations; as these move closer to the miRNA molecule, they will then diffuse away. The resulting equilibrium will be governed by DLVO theory [55 ch 21]. Without an external E field applied, the E field produced by the miRNA molecule gets weakened by the ion cloud. With an external E-field applied, the ion cloud shifts and correspondingly dilutes the effect of the external field, resulting in a final effective negative valence of roughly −18.8*e*^−*a*/*λ*^*^D^*, where *a* is the miRNA radius (≈2.5nm) and λ_D_ is the Debye length of roughly .8nm. This gives an effective charge of ≈-.9, which we conservatively approximate as -1.

### Small tumors depolarize their V*_mem_* as an adaptive behavior to avoid being normalized by surrounding healthy tissue

When a small number of cells with one GRN state are surrounded by a large number of cells with the opposite state, the larger tissue normalizes the smaller one. A new tumor is much smaller than the surrounding healthy tissue, leading to an obvious question: what could prevent it from similarly being normalized – how do small tumors survive?

Tumor cells tend to depolarize [8, 11, 13]; their previous *V*_mem_ of roughly -70mV rises to roughly-20mV. As noted above in Figure 1, a homotypic GJ conducts fully when the two cells it connects have the same *V*_mem_ (so the GJ’s Δ*V*_mem_ is zero). When either of the two cells connected by a GJ changes its *V*_mem_ (so the GJ’s Δ*V*_mem_ is substantially nonzero) GJ conductance falls dramatically.

Given this GJ conductance versus *V*_mem_, the tumor’s electrical behavior seems not only sensible but almost inevitable. By depolarizing (and hence changing its *V*_mem_ to be different from that of healthy tissue), the tumor makes the “border” GJs between the tumor and the healthy tissue become less conductive. This forms a very effective barrier between the tumor and the surrounding healthy tissue, preventing the healthy tissue from normalizing the tumor.

Table 1, using our simulation model, shows how effective the depolarization strategy is. A 6×6×6 cube of tumor cells sits in a 25×25×25 field of healthy cells. Initially, all tumor cells are initialized to the M state ([ZEB] high and [miR200] low); healthy cells are initialized to the opposite (E) state, and GJs are all set to the same base conductivity *G*GJ. We then raise *G*_GJ_, using a binary search, to find the smallest *G*_GJ_ that succeeds in normalizing the tumor. Each column of Table 1 does this under different conditions.

**Table 1:** *tumor strategy affects resistance to normalization.* Each of the three rows is a different amount of GJ underexpression (.3, .5 or .8x that of GJ density in healthy tissue). Each of the four columns is a different tumor-isolation strategy. The numbers are the minimum value of the general system diffusion constant needed for the surrounding tissue to normalize the tumor (units of 10^-15^ meters^2^/sec). We see that while either strategy (depolarizing the tumor cells or underexpressing connexins) is effective, the combination is always most effective at preventing the tumor from being normalized.

|  | Tumor isolation strategy |  |  |  |
| --- | --- | --- | --- | --- |
|  | Neither | Depol. alone | Underexp. alone | Both depol and underexp |
| Underexp=.3 | 1.1 | 38 | 3.3 | 120 |
| Underexp=.5 | “” | “” | 2.1 | 75 |
| Underexp=.8 | “” | “” | 1.3 | 45 |

The first column, labeled “none,” reports the smallest normalizing *G*_GJ_ when the tumor cells are hyperpolarized and GJs are normally expressed. With all cells having the same *V*_mem_, the GJs act as simple conduits for diffusion between cells. The experiment is then repeated with the tumor cells depolarized (and thus more isolated from the healthy tissue) in the second column. We see that this strategy is quite effective, allowing the tumor to withstand much higher GJ density to healthy tissue before being normalized.

But why does a high enough *G*_GJ_ always normalize the tumor, even with tumor depolarization? We will discuss this further when we discuss metastasis; for now, a large enough *G*_GJ_ turns the entire network (both tumor and healthy tissue) into a single effective syncytium. Since the healthy tissue starts with more cells than the tumor does, it will always win such battles.

### Underexpressing GJs: how small tumors further avoid being normalized by surrounding healthy tissue

The next piece of interesting and as yet unexplained data is that not only do tumor cells seem to depolarize, but they also tend to underexpress connexins and thus have a reduced *G*_GJ_.

The “underexpression alone” column of Table 1, again using our simulation model, shows that this behavior is indeed adaptive. It simulates tumor cells underexpressing connexins to levels of 30%, 50% and 80% of that of healthy cells. This leads to both intra-tumor GJs and border GJs having lower conductivity. However, it does not depolarize tumor cells.

Looking at all four columns, it shows that depolarizing the tumor cells provides a strong defense (column 2), that underexpressing GJs provides a somewhat weaker defense (column 3), and that the combination of both strategies is substantially stronger than either one of them alone (column 4). In other words, the two strategies are not only each effective, but are synergistic.

It may not be obvious that underexpressing GJs would be help the tumor resist normalization. The tumor cells most directly under attack by the surrounding healthy tissues are the cells in the outermost layer of the tumor. When the tumor underexpresses GJs, it weakens the connection of the outermost tumor cells to the surrounding healthy tissue, and thus weakens the attack. However, it also weakens the connection of those same outermost tumor cells to the inner core of the tumor, which could support them. So: why is this a net win and not a net loss?

The simulation results, of course, show that it is a net win; for each column that shows underexpression, more severe underexpression (top row) leads to the tumor being more resistant to normalization. But why?

In the 2D picture of Figure 4, the one-cell-wide normalizing healthy-tissue ring, with area *πr*_cell_^2(72-52),^ clearly contains more cells than the helping inner shell of area *πr*_cell_^2(32-12).^ In general, by simple geometry, for a constant shell width 2*r*_cell_, shells of larger radius have larger area. Thus, the outer healthy-tissue shell is always a larger threat than the inner tumor shell is an ally, and reducing GJ density to both is a net win.

**Figure 4:**
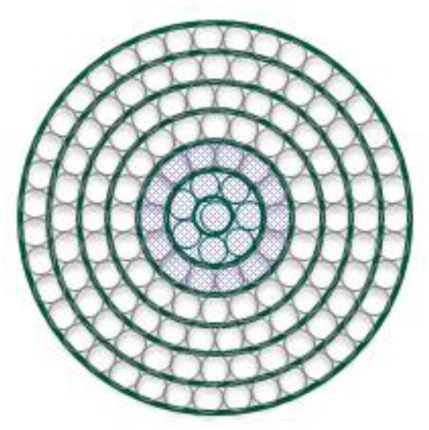
*tumor cells at the border are attacked more than they are defended.* A 2D drawing of a tumor surrounded by healthy cells. All cells are assumed to be in concentric rings (cells drawn with narrow borders and rings drawn with heavier borders); each ring is 1 cell wide. The inner blue cells are tumor cells and the outer cells are healthy. The outermost tumor (blue) shell has outer radius 5r_cell (_with the innermost cell adding r_cell t_o the radius and the other two rings each adding 2r_cell_); it is being attacked by the innermost healthy ring of cells (radius 7r_cell_) and defended by the tumor ring of radius 3r_cell_. The real world is of course 3D (analyzed in the text), but a 2D figure is easier to visualize and draw.

### Proliferating tumors are more robust the larger they are

As noted just above, the cells at the outer edge of a tumor are supported by the tumor cells at its immediate interior, while simultaneously being attacked by the healthy cells just outside the tumor. Simple geometry shows that there are always more attacking cells than defending cells. But are larger tumors more vulnerable to normalization than smaller tumors, or vice versa?

Consider extending the 2D tumor drawing in Figure 4 to 3D: a spherical tumor of radius *r*_tum_, with individual cells being spheres of radius *r*_cell_. The single-cell-thick layer of tumor cells at the exterior of the tumor then has volume 4/3 *π*(*r*_tum_^3-(^*r*_tum_-2*r*_cell_)^3).^ It is attacked by the shell of healthy cells of volume 4/3 *π*((*r*_tum_+2*r*_cell_)^3-*r*3),^ and buttressed by its immediately-innerward tumor shell of volume 4/3 *π*((*r*_tum_-2*r*_cell_)^3-(^*r*_tum_-4*r*_cell_)^3).^

We can measure the direness of the attack by the ratio of the attacking-shell volume to the defending-shell volume, which comes out to 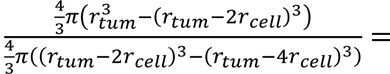 = 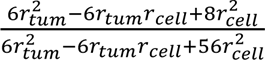. As the tumor (and *r*_tum_) get larger, this ratio is closer to one, and so smaller tumors are thus more vulnerable to attack. This effect has been noted in the literature under laboratory conditions [56, 57].

Table 2 shows this, based on simulation results. The setup is exactly as for column 4 (“both”) of Table 1. However, now each column is for a different tumor size. As predicted, larger tumors are more resistant to being normalized.

**Table 2:** *large tumors better resist normalization.* As in Table 1, each row is a different GJ underexpression. Each column is a different size tumor. The numbers, as before, are the minimum value of the general system diffusion constant needed for the surrounding tissue to normalize the tumor (units of 10^-14^ meters^2^/sec). We see that larger tumors are indeed harder to normalize, as the theory predicts.

|  | Tumor size |  |  |  |  |  |  |  |
| --- | --- | --- | --- | --- | --- | --- | --- | --- |
|  | 1x1 | 2x2 | 3x3 | 4x4 | 5x5 | 6x6 | 7x7 | 8x8 |
| Underexp=.3 | 3.8 | 7.7 | 11 | 15 | 18 | 21 | 23 | 25 |
| Underexp=.5 | 2.2 | 4.6 | 6.9 | 9 | 11 | 13 | 14 | 15 |
| Underexp=.8 | 1.4 | 2.9 | 4.3 | 5.7 | 7 | 8.2 | 9.1 | 9.8 |

### Overexpressing GJs enables large, invasive tumors to create a syncytium

When a tumor gets large enough and is ready to invade nearby tissues, the calculus reverses. It is no longer good enough for the tumor to find a safe corner to grow; it must now convince surrounding cells to work with it. Fortunately for the tumor, it is now large enough to have access to a powerful strategy for doing so – overexpressing GJs. By tightly connecting the tumor cells together, the tumor cells effectively act as a syncytium; one very large cell that now becomes the biggest bully on the block.

Table *3* shows this with our full simulation model. We augment the model with a new parameter *GJ_power*, which describes the amount of GJ overexpression in tumors. If the base conductivity of GJs in healthy tissue is *G*_GJ0_, then the conductivity of GJs within the tumor is *G*_GJ0_\**GJ_power*. The border GJs are then assigned a midway conductivity of *G_GJ_*_0_ ∗ 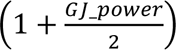 . All tumors are depolarized as before.

**Table 3:** tumor *V_mem_ behavior affects metastasis potential.* Each of the three main columns is a different tumor V_mem_ strategy. Each of the five rows is a different value for the healthy-tissue GJ conductivity (G_GJ0_). The P_min_ values represent the minimum value of the parameter GJ_power that results in successful invasion. The data show that there is no case where invasion is successful without increasing GJ expression inside of the tumor

|  | Tumor metastasis algorithm |  |  |
| --- | --- | --- | --- |
| $G_{GJ0}$ | Depol $P_{\min}$ | Hypol $P_{\min}$ | Neighbor-depol $P_{\min}$ |
| 1e-16 | >100 | 45 | 48 |
| 3e-16 | 95 | 14 | 17 |
| 2.6e-15 | 45 | 4.1 | 6.0 |
| 7.8e-15 | 21 | 1.5 | 2.4 |
| 2.3e-14 | 11 | 3.3 | 1.4 |

Each row of the table shows a different value for *G*_GJ0_. The column labeled “Depol P_min_” shows the minimum value of *GJ_power* for which the tumor successfully flips all cells in the surrounding tissue. E.g., for *G*_GJ0_=7.8×10^-15,^ the GJs in the tumor must be roughly 21x larger than that for successful invasion.

Let’s try to understand these numbers a bit better. For the upper rows of the table, *G*_GJ0_ is “unreasonably” small, such that no matter how strong the tumor is, there is no way to flip a healthy cell without greatly increasing the conductivity of border GJs, which, as noted above, are scaled up by .5*GJ_power*.

Values of *G*_GJ0_ larger than the 2.3×10^-14 (^not shown) are “unreasonably” large; they turn the healthy tissue into a syncytium, and one that is larger than the tumor. They would thus not enable metastasis. The intermediate table rows are the interesting values, where GJ overexpression stays reasonably constant. The table columns labeled “neighbor-depol P_min_” and “hypol P_min_” will be discussed shortly; they represent tumor strategies that use *V*_mem_ changes to aid GJ overexpression and further facilitate metastasis.

If overexpression is so effective, why didn’t the tumor use it from the beginning? Because aggregating a large tumor into an effective syncytium is only effective if the tumor is, in fact, large. Table 1 showed this; with a large enough *G*_GJ_, the surrounding healthy tissue overwhelmed and normalized the small amount of tumor tissue.

### Manipulating V*_mem_* allows the syncytium to connect to healthy tissue and corrupt it

Overexpression of GJs within the tumor leads to the tumor acting as a syncytium. However, that is of limited use unless the tumor also has a low-resistance path to attack healthy tissue. When the cells in a tumor depolarize, the border GJs have increased resistance, which hinders the tumor from altering the state of nearby healthy cells. This is why it required such high values of GJ_power to induce metastasis in the “Depol P_min_” column of Table *3*.

How can an invasive tumor increase GJ conductivity at its borders? The simple fact of connexin overexpression within the tumor may also increase the tumor’s GJ connectivity to nearby healthy cells. Remember that a GJ forms when two adjacent cells, each expressing connexins and forming a hemichannel, dock to each other. Though the process is not fully understood [22], it seems reasonable that the more connexin the tumor expresses and transports to the cell membrane, the more hemichannels it will form and the more of these hemichannels are likely to dock with healthy cells.

A powerful strategy is simply for the tumor to now hyperpolarize its cells again. Remember (Figure 1) that homotypic GJs have their highest conductance when Δ*V*_mem_ across them is near zero, and conduct less as |Δ*V*_mem_| gets larger. Depolarizing the tumor was an effective GJ isolation tactic because, with the surrounding healthy tissue hyperpolarized, it largely turned the border GJs (the GJs between the tumor and surrounding tissue) off. By the same token, re-hyperpolarizing the tumor sets Δ*V*_mem_ across these border GJs near zero again, causing the border GJs to fully turn on and thus reconnecting the tumor to healthy tissue.

While this does render the tumor vulnerable to attack by nearby healthy tissue, the tumor is now large enough to withstand such attacks. More to the point, it increases the tumor’s ability to invade and metastasize. The “hypol P_min_” column of Table *3* shows detailed simulation results with this strategy. Notably, successful invasion now requires substantially less GJ overexpression.

However, evidence as to the actual *V*_mem_ of invasive tumors is quite mixed. While some work has indeed found hyperpolarization [15], the majority of other work has found either continued depolarization [12, 13, 20], slowly moving *V*_mem_ waves linked to changes in cell volume to promote motility [12, 13, 20], or negative-going action-potential spikes from a depolarized base level [58].

Can a tumor connect itself tightly to neighboring healthy tissue without hyperpolarizing the tumor cells? Surprisingly, a clever strategy by the tumor can indeed do so. While the tumor remains depolarized, it could recruit nearby healthy cells to *also* depolarize. Those nearby healthy cells now share the same *V*_mem_ as the tumor and have a different *V*_mem_ from the mass of surrounding healthy cells; they thus become tightly connected to the tumor (ensuring that the tumor can flip their state) and largely disconnected from the support of their healthy surrounding tissue.

The “neighbor-depol P_min_” column of Table *3* shows detailed simulation results with this strategy. This strategy tends to be slightly more effective than simple hyperpolarization.

### Our intuitive simulation model

All simulation results above use the detailed EMT model. However, as noted, we do not claim to know precisely what constitutes tumor state; different tumors almost certainly store state in different ways. It thus seems useful to augment our reasonably detailed model with a more abstract and simpler model that will apply equally well to other pathways, as well as give strong intuition and allow analytical analysis.

In this model, we linearize a cell’s detailed model at a particular steady-state (SS) operating point, creating a small-signal linear model that allows easy analysis. This analysis is of course less valid the further the system strays from the given operating point.

Start with generation and decay of a generic miRNA *mR* in a single cell *A*:

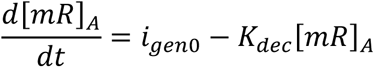

*I*_gen0_ is the generation rate of *mR* in cell *A*, in moles/m^3s,^ and is simply the Hill-model value given, say, a logical “zero” state of cell *A* (a logical “one” state would have generation rate *i*_gen1)_. *K*_dec i_s the decay rate for *mR*, in s^-1 (^assuming for simplicity that the same linear degradation rate applies to *mR* in its free and bound forms).

This reaches a SS value of [*mR*]_A_=*i*_gen0_/*K*_dec_. Next, add in an external current drain *i*_exit,A_ (which will model miRNA exiting the cell through a GJ)

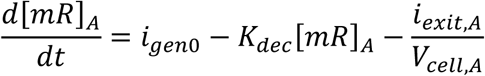

*I*_exit,A_ has units mol/s and *V*_cell,A_ is the volume of cell *A*. The new SS value is [*mR*]*_A_* = 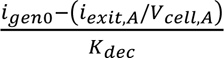. We define *mR*_UL0_≡*i*_gen0_/*K*_dec_ (mol/m^3)^ and *G*_eq,A_≡*V*_cell,A_*K*_dec_ (m^3/s^) to give

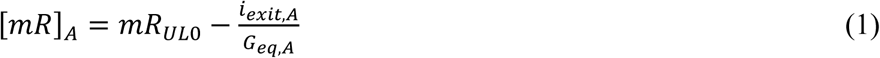

Intuitively, *mR*_UL0_ is the SS concentration of *mR* in a “0” cell under “unloaded” conditions (where *i*_exit,A_=0). *G*_eq,A_ is the cell’s resilience to an external current draw; the higher *G*_eq,A_ is, the less *i*_exit,A_ affects SS [*mR*]_A_. When an external system pulls *i*_gen0_*V*_cell,A_ mol/s of *mR* out of the cell (i.e., the entire generation current, in mol/s), then the 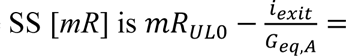 = 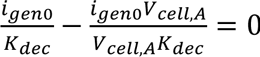.

If the only characteristic of cell *A* that we care about is its SS behavior at this single port, then equation (1) fully captures that behavior, and we have no need to go back to the original differential equation. We can use this fact to more easily analyze the two cells interconnected by a GJ in Figure 3a.

When two cells with different [*mR*] suddenly communicate via a GJ, each affects the other’s [*mR*]. The system of two cells and one GJ will obey the black-box equations (assuming cell *A* in the 0 state and cell *B* in the 1 state)

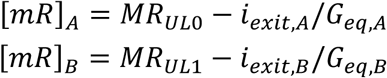

Since *i*_exit,A_ and *i*_exit,B_ both enter the same GJ, then at SS they must obey *i*_exit,A_=-*i*_exit,B_. For simplicity, we will assume drift currents are negligeable. Then Fick’s Law for the GJ is *i*_exit,A_=*G*_GJ_ ([*mR*]_A_-[*mR*]_B_). This quickly yields (defining *G*_rel,A_≡*G*_eq,A_/*G*_GJ_ and *G*_rel,B≡_*G*_eq,B_/*G*_GJ_),

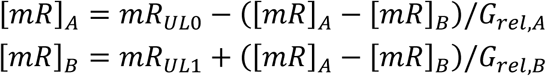

or

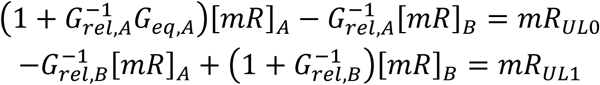

or

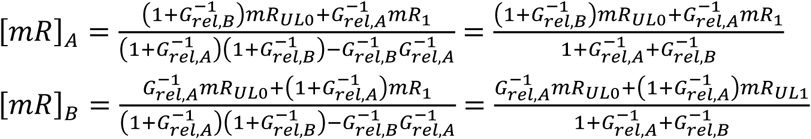

or the simple weighted sums

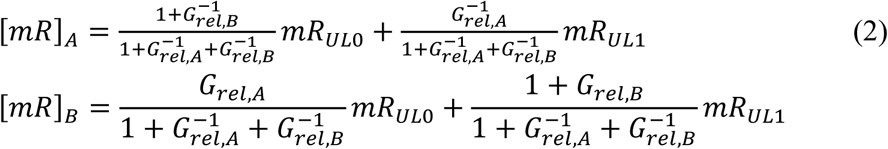

These very simple equations will serve as intuition for the effect of changing GJ density on communication between cells, and will explain both the isolation and the syncytium behaviors noted in this paper.

### Using the intuitive model

Let’s use our simplified model to better understand the experiments summarized in Table 1. The bistable GRN in each cell has a metastable point, which is somewhere between *mR*_UL0_ and *mR*_UL1;_ if the model predicts that interconnecting the cells would move a cell’s [*mR*] past the metastable point, then that cell’s state will flip. But Figure 3*b* shows clearly that the lower the value of the GJ conductance *G*_GJ,_ the more each cell retains its original value, staying far from the metastable point and thus holding state. Intuitively, then, the lower *G*_GJ c_aused by tumor depolarization does indeed isolate the tumor.

Equation (2) has the form of weighted sums, with 1/*G*_rel,A a_nd 1+1/*G*_rel,B b_eing the weights to determine whether the final value of each cell is closer to *mR*_UL0 o_r *mR*_UL1._ Remembering that *G*_rel,A≡_*G*_eq,A_/*G*_GJ_ = *V*_cell,A_*K*_dec_/*G*_GJ_. Intuitively, very high *G*GJ gives equal weights and forces both cells to roughly (*mR*UL0 or *mR*_UL1_)/2; very low *G*_GJ_ allows each cell to maintain its [*mR*].

^Fig^ure ^3^b shows that with a high enough *G*_GJ_, all cells will eventually reach the same [*mR*]. But will this [*mR*] correspond to a tumor state or a healthy state? Why, when we raised *G*_GJ_ high enough, did all of the isolation experiments show tumor normalization?

If one cell is larger than the other, equation (2) predicts that our two-cell system will have the final [*mR*] closer to the initial value of the larger cell (again, with cell size reflecting into *G*_rel_). Figure 5a shows the results of cell *B* being 1.5x larger than cell *A*. But we would not expect drastic cell-size differences between tumor cells and healthy cells as much as simply differences in the number of cells. In fact, a large enough *G*_GJ_ turns the interconnected cells into a functional syncytium.

**Figure 5:**
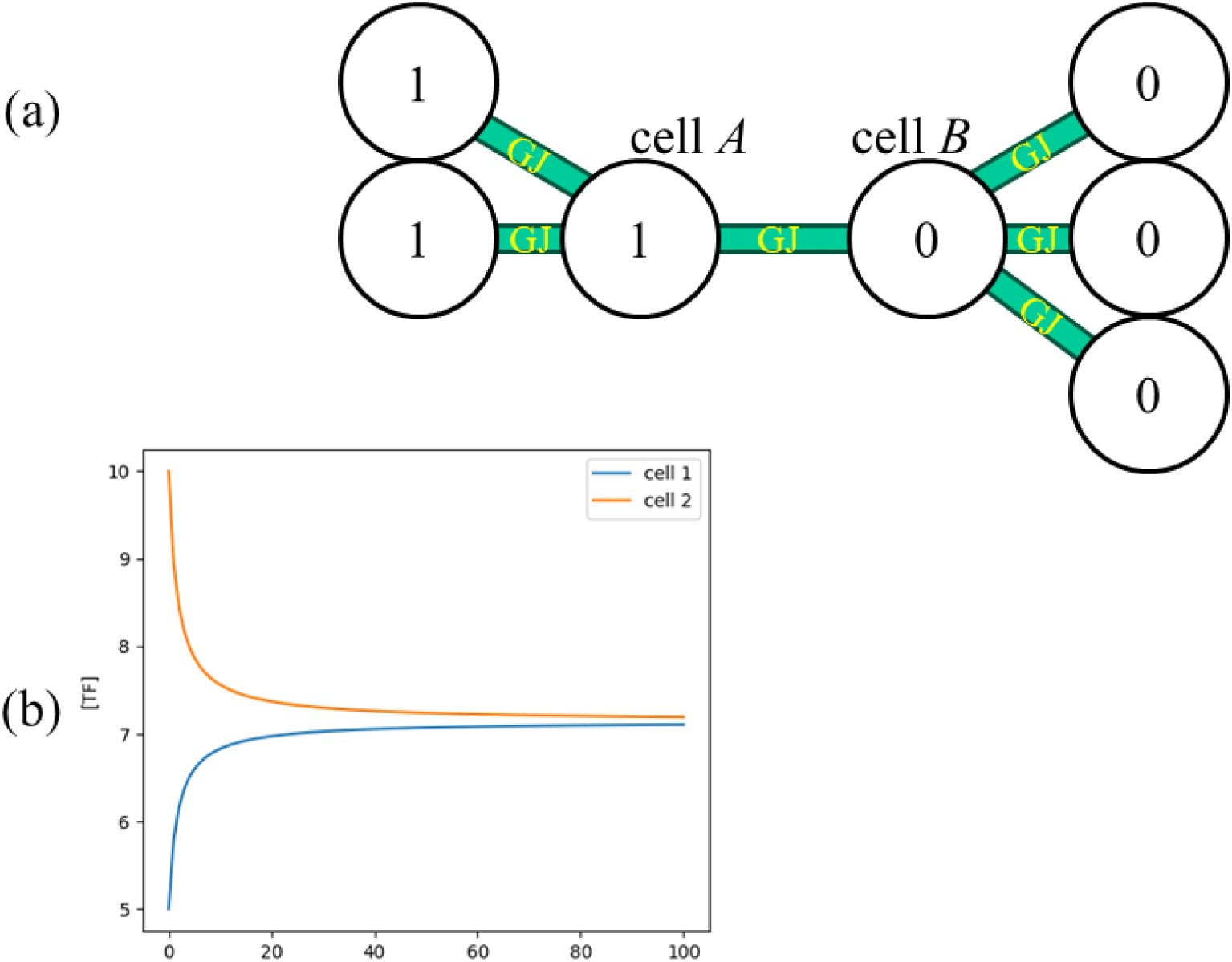
*multiple cells connected by GJs.* (a) Two cells connected by GJs, now with five backup cells. (b) Steady-state [mR_T_] in each cell. Compared to Figure 3, we now model cell B as being 1.33x larger than cell A, resulting in the final value being closer to the cell B initial value (7.15 rather than 7.5)

A simple analogy is that of batteries in parallel. If you connect a 12V car battery in parallel with a 1.5V AAA battery, then (after a small amount of sparks!) the voltage of the common terminal will be close to 12V. Why is the AAA battery forced to take the voltage of the 12V battery, rather than the net voltage being, e.g., the average (12+1.5)/2=6.75V? Because the 12V battery has a vastly-lower internal resistance, due to its internal structure of a large number of smaller cells in parallel.

We will shortly formalize this concept. For now, as we raised *G*_GJ_ in our isolation simulation, we eventually turned the tumor into one syncytium and the healthy tissue into a second syncytium. As there were more healthy cells than tumor cells in the simulation, the final [*mR*] was healthy.

Equation (2) and Figure 5b show that essentially the larger cell wins. The more the healthy tissue is larger than a small tumor, the greater the certainty that it normalizes the tumor rather than vice versa. This explains why there always is a minimum *G*_GJ_ at which normalization occurs in Table 1.

Now let’s look at GJ overexpression during metastasis. When a tumor overexpresses GJs to more tightly interconnect the tumor cells, it turns itself into a syncytium – a giant cell that, like the 12V car battery, can overwhelm any normal-sized healthy cell. However, the tumor is surrounded by healthy cells over its entire border, which is a more challenging environment.

Figure 6 illustrates a large tumor, shown as a 2-D hexagonal array of light-red cells. The exterior ring of tumor cells is surrounded by healthy tissue, of which only the inner ring is shown. Metabolites pass across the border as the cells try to flip each other’s states. We would like to compute who wins the tug of war.

**Figure 6:**
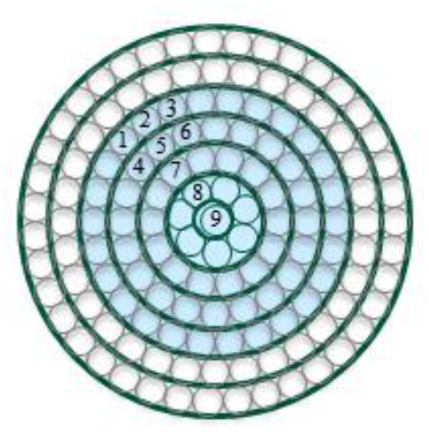
*an external cell of a syncytium is best analyzed as a linear slice.* This figure follows the same notation as Figure 4, showing a group of tumor cells (colored light blue) surrounded by healthy tissue. Assume that GJs connect all neighboring cells. A representative cell (#2) is attacked by healthy cells. It may seem that cell #2’s tumor state is supported cells #4, 5 and 6; however, #4 is simultaneously being attacked by #1, and #6 by #3. Thus, symmetry considerations imply that cell #2 is only supported by the straight line (cells #5,7,8,9) to the tumor core. A pie slice would be a more accurate model, but more difficult to analyze.

With all the exterior tumor cells simultaneously under attack, any given exterior tumor cell does not have the entire tumor core for its exclusive support. Rather, it can only depend on the single pie slice from itself to the tumor’s center. For simplicity, we will model this slice as a straight chain of cells.

We can apply our simple model recursively to analyze this connected chain. Figure 7a recaps our black-box equivalent of a single cell. The black box considers only two variables: the output current draw (mol/s) of *mR*, and the resulting 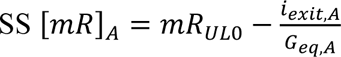, where *i*_exit,A_ is the black box input (mol/s), *mR*_UL0_ is the steady state concentration [*mR*] at *i*_exit,A_=0, and *G_eq_*_,*A*_ models the slope of the input-output relationship.

**Figure 7:**
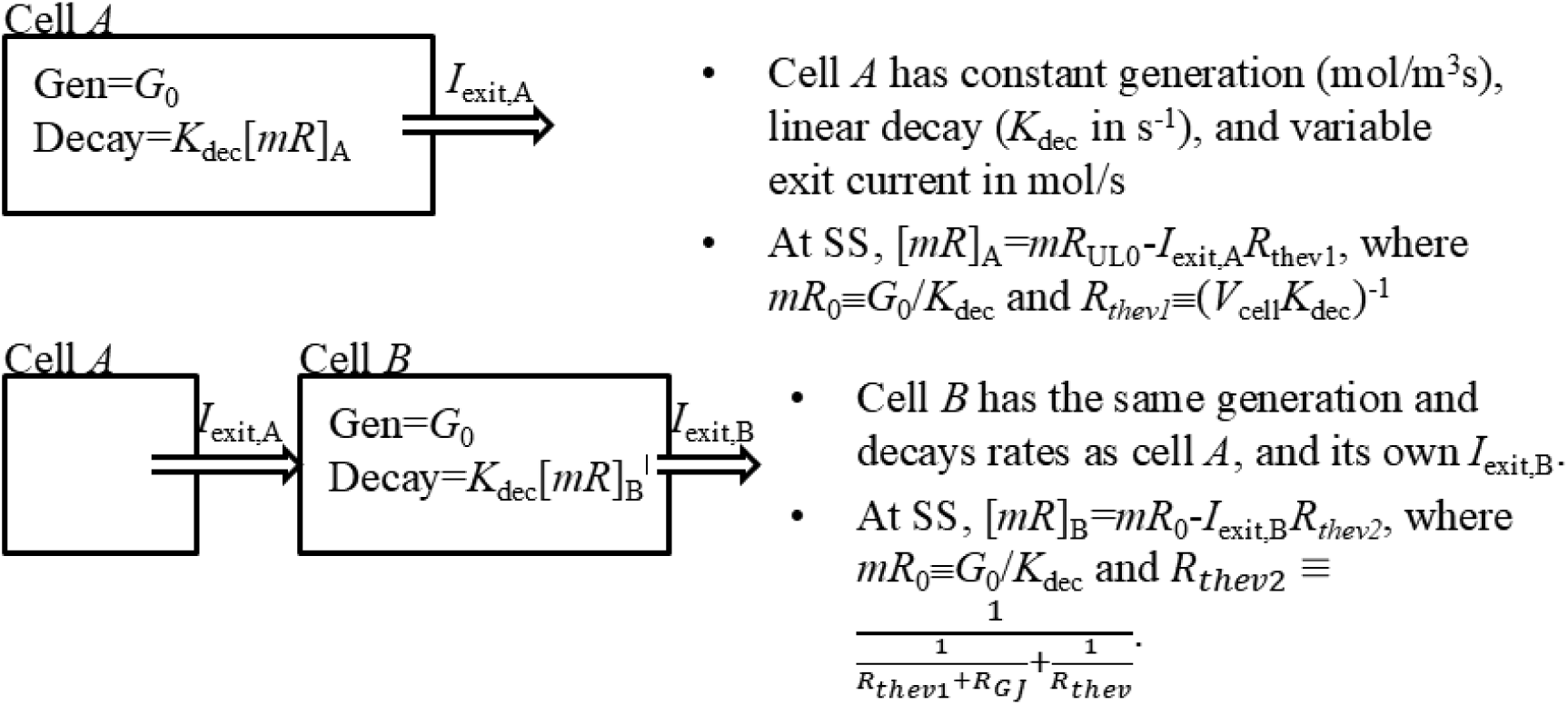
*a recursion step helps analyze the syncytium.* The top drawing shows one cell with generation, decay and an exit current. At SS, it can be modeled as a black box with [mR]_A_=mR_0_-I_exit,A_R_thev1_. This is merely our basic model again. The bottom drawing shows two cells. Rather than analyze the full system of coupled differential equations, we get the SS solution by modeling cell A as a black box. This procedure can be carried out repetitively to arrive at a syncytium model.

The appendix derives the recursion step, (Figure 7b), moving from a chain of *n* cells to a chain of *n*+1 cells, as [*mR*]*_n_*_+1_ = *mR_UL_*_0_ − *i_exit_*_,*n*+1_/*G_eq_*_,*n*+1_, where *G_eq_*_,*n*+1_ ≡ 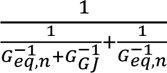.

Since the black-box model for this entire system has exactly the same form as that for cell *A*, the process can easily be carried out for longer and longer chains of cells.

Figure 8 shows the results for successively longer cell chains. Each plotted line is the black-box model for a chain of cells. In this model, *i*_exit_=0 describes the undisturbed system, with no external cells trying to move *MR* into or out of the system. Since each individual cell, if undisturbed, would reach the same steady state concentration *mR*_UL0_, so do all of the chains; hence the common intercept for all chains of *i*_exit_=0, [*mR*]=*mR*_UL0_.

**Figure 8:**
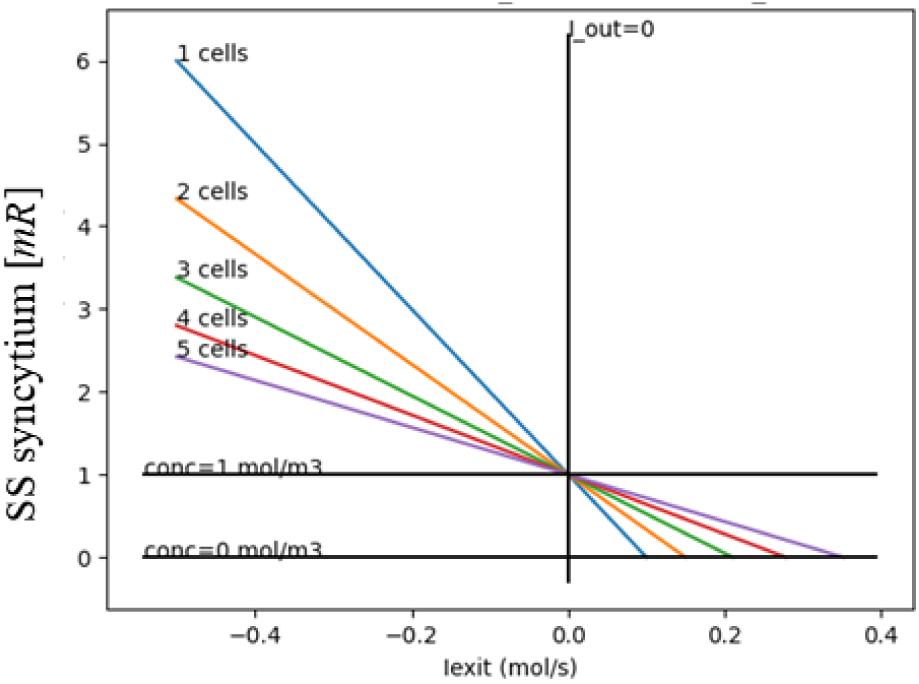
*larger syncytiums proselytize better.* Each colored line plots the response of a syncytium of a different size (represented by a straight chain of cells of that length). The x axis shows how much mR the syncytium is supplying to the external tissue; the y axis shows the resultant syncytium [mR]. All syncytiums nominally have [mR]=1 mol/m^3.^ The syncytiums with more cells can supply more mR to the healthy tissue while seeing their own [mR] be disturbed less.

What differs is that adding more and more cells to the chain (i.e., making a syncytium from a larger and larger tumor) reduces the slope of the graphs; it means that an external source or sink of mR molecules makes less and less difference in [*mR*] at the tumor border. This precisely means that the tumor will be more and more dominant at enforcing its [*mR*] and flipping the state of neighboring healthy cells rather than being flipped itself.

This high-level technique for analyzing steady-state metabolite levels is very reminiscent of Thevenin equivalent circuits from electrical engineering. Our *MR*_UL0_ is acting as an open-circuit voltage and our *G_eq_* acts as a Norton-equivalent conductance. The analogy could be taken further; cell volume could be treated as a capacitor, and all of our differential-equation algebra could have been replaced by simple circuit analysis, with current sources corresponding to metabolite generation, and resistors corresponding to degradation and to GJs. We chose not to present it this way to avoid confusion between the equivalent voltage sources that represent concentration in mol/m^3^ and the actual voltage sources for *V*_mem_.

### Testable predictions

The main strength of our hypothesis is that it explains a large number of existing data points for *V*_mem_ and for GJs in one framework. However, we also make predictions for experiments that have not yet been run:

- If a tumor stays depolarized during invasion, its *V*_mem_ depolarization will extend slightly past the tumor edge.
- If we can build tumors of different shapes, then cubic tumors should invade better than oblong ones of similar volume, and also resist normalization better.
- Larger tumors will more successfully invade than smaller tumors, up to a certain size. If these experiments confirm our hypothesis, it may lead to therapies that use both *V*_mem_ and connexin misexpression as a joint marker of which cells to attack. It may also help to focus future research on cancer messaging through GJs and on better means of controlling tumor *V*_mem_.

## Acknowledgements

We gratefully acknowledge support for this work provided through a sponsored research agreement with Astonishing Labs.

## Conflicts of Interest

This research was funded at Tufts under a Sponsored Research Agreement with Astonishing Labs. M.L. is a co-founder and shareholder of Astonishing Labs.

Astonishing Labs has certain rights to any inventions associated with this research.

## Data availability

All data and code are publicly available on Gitlab.

## Discussion

We have restricted our analysis to homotypic GJs. However, there is evidence of heterotypic GJs involved in intravasation at the tumor border, both in mouse melanoma [30] and in human squamous-cell lung carcinomas [31]. One important difference between heterotypic and homotypic GJs is that while homotypic GJs conduct best at Δ*V*_mem_=0 (Figure 1), heterotypic GJs typically have their conduction curve shifted [24]. E.g., a heterotypic Cx43/Cx45 GJ conducts best when the Cx45 cell is about 25mv higher than the Cx43 cell [59]. If invasive tumors really do create heterotypic GJs at their border, it means that staying depolarized while healthy tissue is hyperpolarized may become a very effective strategy for rendering the border GJs highly conductive.

The concept of bistable cells trying to flip the state of nearby cells is not new; it is well established as a strategy to reliably read embryonic morphogen gradients. Both Drosophila and the vertebrate neural cord, two of the most widely studied models of embryonic development, use this technique [60–62] to interpret morphogen gradients and thus divide a field of cells into a small number of distinct segments (the *positional information* theory of morphogenesis [63, 64]). The cellular environment is noisy, and some small number of cells may misread the gradient. Both model organisms thus use a bistable GRN in each cell to have the majority of correct cells flip the occasional incorrect cell, exactly as we imagine happening in tumor cells.

Drosophila and the vertebrate neural cord both use diffusible protein transcription factors as messengers. The early Drosophila embryo is a syncytium, and proteins can thus diffuse easily between nuclei. In the vertebrate neural tube, however, the cells are fully separate; thus, long-range signaling within the neural tube relies on secreted proteins [60] received by transmembrane receptors and transduced by intracellular signaling pathways. Two very different species evolved very different mechanisms for communicating between cells; but the concept of using bistable GRN nodes, where a majority of cells flip the state of errant minority cells, evolved in both. Furthermore, there is evidence [65] that miRNA is the messenger for similar functionality to decode rostral-caudal position along the spinal cord.

The ability of a majority of correctly-decoding cells to “convince” the minority of incorrect cells embedded in them to flip their state value is essentially a form of entrainment, where the majority cells normalize the minority cells into the majority state. This is a case where normal morphogenesis uses the technique, and cancer has co-opted it – but has also combined it with GJ overexpression to become more potent.

Interestingly, some animal models of cancer also seem to fit this paradigm. [66] induced melanomas in Xenopus laevis embryos by depolarizing a small number of “instructor” cells. The depolarization was not 100% effective; it only induced melanoma in some embryos, but not all. However, any single embryo was either fully converted to the invasive phenotype or not converted at all. There were no DNA changes in these Xenopus experiments; the only cellular changes were in non-DNA state. In other words, whatever state changes occurred in the instructor cells were propagated to all melanocytes in the embryo, similar to what we observed in our metastasis simulations.

It may seem surprising that a healthy cell could be converted to a cancerous phenotype without DNA changes. However, other similar examples have been noted [67]. Furthermore, the cells whose *V*_mem_ were manipulated in [66] are *not* the cells that became metastatic; this shows the importance of the microenvironment in determining cancer phenotypes [7]. And since GJ-related mutations are *not* commonly seen in tumors [16], it is likely that the changes in GJ expression are caused by GRN state.

The strategy of underexpressing GJs as a means of isolating tissue from its surroundings is not restricted to cancer. Planaria are a type of flatworm that are able to regenerate their entire body from small fragments. Experiments [68] have submerged planaria in a barium chloride solution, which causes the worm’s head to degenerate. Amazingly, the worm figures out how to regrow its head while still submerged in the BaCl. RNA sequencing of the regenerated worm showed that it was underexpressing GJs; one explanation is that this enables the majority of the regenerating worm to isolate itself from the toxin. I.e., the concept of isolation was originally used by healthy tissue, and has been co-opted by cancer.

As noted in our Introduction, the evidence in [30–32] has been used to hypothesize [18] that the tumor uses GJs to send metabolites to alter the neighboring healthy tissue and thus ease invasion. In general, the use of chemical messages to influence a tumor’s surroundings is well established. Specifically, messaging between tumor cells and cancer associated fibroblasts, alters fibroblasts subtype, thus helping to structure the tumor micro-environment. (reviewed in [69]).

## Conclusion

The electrical behavior of tumors, while well known, has resisted a simple, unifying explanation. We have tried in this paper to provide one by viewing it from a standpoint of electrical communication through GJs.

When a small tumor depolarizes itself, the depolarization automatically disconnects the tumor from its surrounding healthy tissue, avoiding communication that could help the healthy tissue normalize the tumor. By underexpressing GJs, a small tumor further acts to prevent communication to the outside world, and hence prevents itself from being normalized.

When a tumor becomes large, it can now employ a new strategy to help it invade surrounding tissues. By overexpressing GJs, it can act as a syncytium, essentially giving it the ability to act as a single extremely dominant cell that can easily proselytize its neighbors. It can then re-hyperpolarize itself to make a communication channel to enhance this proselytizing; or it can remain depolarized and instead alter the *V*_mem_ of nearby healthy cells to, again, create that communication channel. The large size of a tumor increases its volume-to-surface-area ratio, further enhancing its ability to dominate communication.

## Figures

- Figure 1: the conductance versus voltage for a GJ
- Figure 2: two cells connected by a GJ, with a graph showing their behavior
- Figure 3: two cells connected by a GJ, but with numerous cells behind them
- Figure 4: rings of tumor and healthy cells showing why larger tumors survive
- Table 1: isolation results for GJs, underexpression and both
- Table 2: results showing larger tumors survive better
- Table *3*: results showing metastasis success rates for different tumor strategies
- Figure 5: multiple cells connected by GJs
- Figure 6: the pie-slice assumption behind our metastasis model
- Figure 7: the recursive model showing why invasive tumors overexpress GJs
- Figure 8: graph of invasive ability vs. tumor size

## Appendix: syncytium calculations

Consider Figure 7. The top drawing merely recaps our existing black-box model of a single cell.

Next (bottom drawing), connect cell *A* to a cell *B* via a GJ. The GJ has the usual *G_GJ_* = 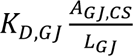 in m^3/s^; cell *B* has the same internal generation rate *i*_gen0_ and volume *V*_cell_ as cell

*A*; it is driven by *I*_exit,A_ via the GJ, and has its own exit rate *I*_exit,B_. We would like to put together a similar model for this system, where *I*_exit,B_ predicts [*mR*]B. So given *I*_exit,B_, what is [*mR*]_B_?

Let’s first compute *I*_exit,A_ given [*mR*]_B_. From equation (1) above, we have [*mR*]_A_=*mR*_UL0_-*i*_exit,A_*/G*_eq,A_. Now, all the flow exiting cell *A* enters the GJ; so the GJ flow into cell *B*, in mol/s, is by definition *i*_exit,A_. But by Fick’s Law, the GJ flow in mol/s is also given by flow = *i*_exit,A_ = *G*_GJ_ ([*mR*]_A_-[*mR*]_B_). Inverting this gives 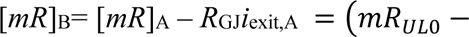 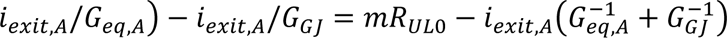. Rearranging again gives 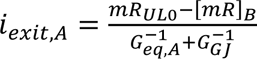.

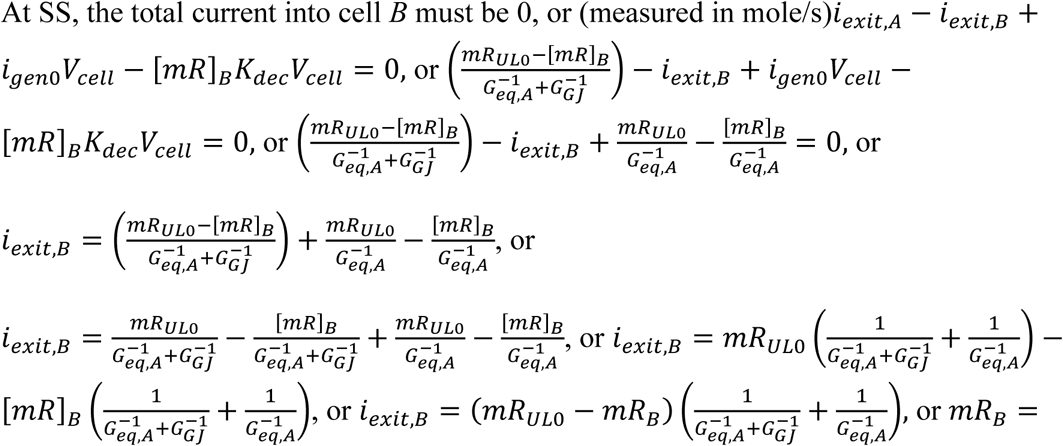

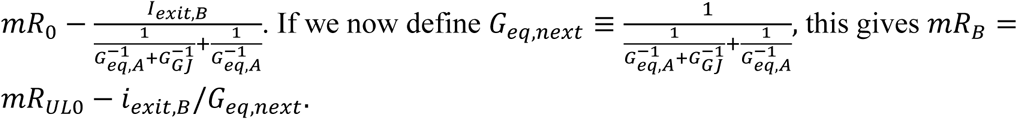

- The equivalent circuit always has the form [*mR*]*_n_*_+1_ = *mR_UL_*_0_ − *i_exit_*_,*n*+1_/*G_eq_*_,*n*+1_, where 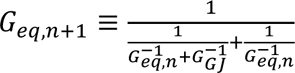

